# Untangling biological factors influencing trajectory inference from single cell data

**DOI:** 10.1101/2020.02.11.942102

**Authors:** Mohammed Charrout, Marcel J.T. Reinders, Ahmed Mahfouz

**Affiliations:** Delft Bioinformatics Lab, Delft University of Technology, Delft 2628 XE, The Netherlands; Leiden Computational Biology Center, Leiden University Medical Center, Leiden 2333 ZC, The Netherlands; Department of Human Genetics, Leiden University Medical Center, Leiden 2333 ZC, The Netherlands

**Keywords:** single cell, pseudotime, factor analysis, trajectory inference

## Abstract

Advances in single-cell RNA sequencing over the past decade has shifted the discussion of cell identity towards the transcriptional state of the cell. While the incredible resolution provided by single-cell RNA sequencing has led to great advances in unravelling tissue heterogeneity and inferring cell differentiation dynamics, it raises the question of which sources of variation are important for determining cellular identity. Here we show that confounding biological sources of variation, most notably the cell cycle, can distort the inference of differentiation trajectories. We show that by factorizing single cell data into distinct sources of variation, we can select a relevant set of factors that constitute the core regulators for trajetory inference, while filtering out confounding sources of variation (e.g. cell cycle) which can perturb the inferred trajectory. Script are available publicly on https://github.com/mochar/cell_variation.

**Significance Statement:** Pseudotime inference is a bioinformatics tool used to characterize and understand the role and activity of genes involved in cell differentiation. To achieve this, the level of expression of thousands of genes are simultaneously used to order cells along a developmental axis. However, this may result in distorted trajectories as many genes are not necessary involved in cell differentiation, and might even provide the pseudotime inference tool with conflicting (confounding) information. Here we present a methodology for improving inference of the differentiation trajectories by restricting it to a small set of genes assumed to regulate cell differentiation.

---

Single cell RNA-sequencing enables quantitative gene expression profiling of individual cells. From an RNA viewpoint, these cells live in a high-dimensional space defined by the expression of their genes. A critical step when analyzing such data is the identification of cells in order to find and label the cell types present in the data. This is often achieved by grouping together cells with similar expression profiles by applying a clustering method. The resulting cell clusters are thus separated from one another by a set of genes uniquely expressed or silenced in a subset of clusters. These so-called marker genes can then be used for identification by cross-referencing with known marker genes or marker genes found in other studies. This clustering-based approach for cell identification relies on the general presumption that the measured expression levels are reflective of the cell’s identity, which may be violated due to shared transcriptional programs between two or more types. Large variations within cell type clusters due to many exclusive programs may also pose a problem as it can become hard to discern between cell types and cell states (1). More generally, sources of variation that contribute significantly to the cell-cell distances in gene space, yet do not reflect the cell type, can be detrimental to the identification task. These can vary from small transient changes e.g. cell communication, up to complex shifts in the cell’s regulatory state such as the cell cycle, which has been reported to contribute a substantial portion of the gene expression variance (2). Moreover, cell identification is often preceded by a gene filtering step whereby genes with low variance (and therefore little information) are discarded to ease the computational burden in downstream analysis. Depending on the normalization and filtering criteria used, gene filtering can lead to a lower dimensional space that further amplifies unwanted variabilities. It becomes clear then that identifying and filtering out unwanted biological sources of variation can serve an important step in cell identification.

The immediate question to ask is how to identify what genes are necessary and what genes need to be left out when carrying out such analysis, which leads into a broader discussion of the definition of a cell type. Classically, cells were characterized using a combination of morphology, lineage, location and overall cell function. (1, 3) However it has long been demonstrated that terminally differentiated cells can convert into other selected cell types by overexpression of key regulators (4–6). It has therefore been argued that cell types can be identified by the expression of a unique combination of transcription factors that make up the core regulatory complex, which is preserved along all states of the cell (1, 7). Successfully identifying these core regulators allows one to differentiate between clusters of cell types and clusters of cell states, as the latter would share core regulators. Furthermore, focusing only on this stable set of differentially expressed genes relieves us from determining what transcriptional programs relate to the identity of a cell. In developmental systems, before complete cell maturity is reached, developing cells have been shown to undergo a series of discrete metastable states here referred to as differentiation checkpoints (7–9). By relaxing the aforementioned definition of regulatory complexes to also include differentiation checkpoints, an analogous approach can be used to identify types when dealing with continuous cell transitions.

We focus on the problem of pseudotime inference where the aim is to order developing cells along a “pseudotime” axis based on their transcriptional similarities. These similarities should therefore strictly reflect differences between cell types as they progress through the differentiation trajectory. The majority of pseudotime inference tools rely on the existence of a continuous manifold that reflects this trajectory such that a 1-dimensional curve or graph can be fit (10). Confounding biological sources of variation (such as the cell cycle) can therefore perturb the inferred trajectory. We therefore hypothesize that by factorizing the matrix into distinct sources of variation, a relevant set of factors that constitute the core regulatory complexes can be selected for improving trajectory analysis.

## Results

### Overview

A summary of our approach is visualized in figure 1. First, the count matrix is factorized into a set amount of factors, each representing a transcriptional program. For this, scHPF (11) was used, which assigns for each factor a score per gene that quantifies the contribution of that gene to the factor. Similarly, cells are assigned a score based on how active the factor is in the cell. Scores concentrated on a select subpopulation of cells and a small subset of genes indicate specialized processes, whilst factors with a more uniform score distribution indicate broader processes active in many cells. The number of factors is selected based on the presumed number of major and intermediate cell types. However, by increasing this number of factors, the resolution can be increased from broad subpopulations to highly specific cell states. Next, the factors representing the cell types and differentiation checkpoints are selected. This is a manual process done by a combination of known marker genes, Gene Ontology annotations, the number of highly variant genes, and preservation of the factors across different runs. Next, the top 10 factor genes are passed to Ouija (9), a pseudotime inference method that models gene expression directly as either switch-like, where the gene is activated or repressed at some point during differentiation, or transient, when expression is only active for a short period of time. The direct modelling of a small set of marker genes allows for more interpretable inference and is therefore more in line with our biologically motivated approach. However, Ouija is limited to linear, non-branching data but can be used nonetheless by repeating the process for each sequence of progenitor to mature cell type factors.

**Fig. 1.**
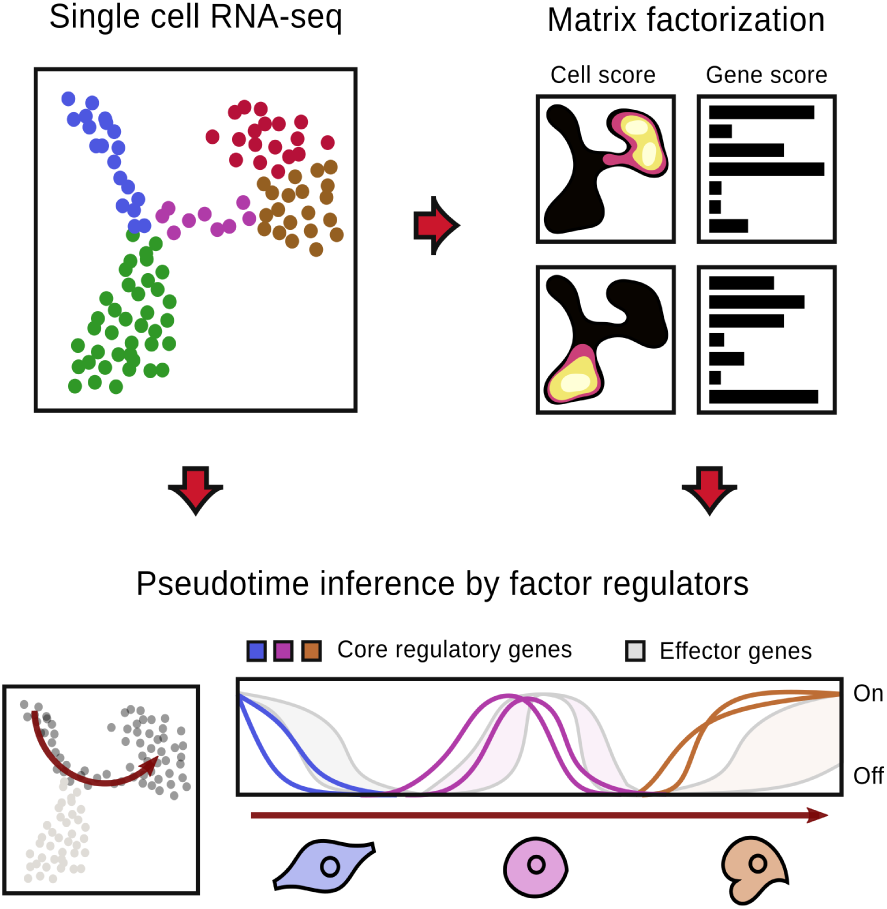
A graphical overview of the methodology. A single cell RNA-seq dataset of developing cells is first decomposed into a set of factors using matrix factorization. Each factor captures a different source of variation, a subset of them of which constitue the core regulatory genes driving the differentiation. A gene-based modelling approach is then used to find the unobstructed trajectory using only the regulatory genes of the differentiation factors.

### Cycling progenitor cells lead to spurious embeddings

As a first demonstration of our methodology, we study radial glia cells that are known progenitors of glia cells, the astrocytes and oligodendrocytes, as well as granule and pyramidal neurons in the developing hippocampus (13). When activated from their quiescent state, radial glia differentiate into neuronal intermediate progenitor cells (nIPC) and undergo continuous cell division during the development period, which is reflected strongly in the transcriptomic profile of these cells. To show to what extent this cell state can affect the analysis of the continuous embedding, the cells of the developing mouse hippocampus from La Manno et al. (12) were reanalyzed, focusing on the subset affected by the cell cycle by excluding the neuron specification branch. Figure 2A shows the UMAP embedding based on the top 3000 most variant genes, annotated by cell types as identified by the data source. The effect of the cell cycle was analyzed by first identifying a set of three factors associated with the cell cycle (figure 2E), and subsequently reconstructing the embedding twice, with and without the cell cycle factors (figure 2B and C respectively). A significant difference in the resulting embedding can be observed, which is further quantified by the jaccard distances of the cell neighbours visualized for both embeddings in figure 2D. In all embeddings, the radial glia progenitor cells, as well as the astrocytes, neuroblasts, and oligodendrocytes precursor cells (OPC) are clearly separated, with developing cells forming a bridge between all four clusters. Of note are the nIPC cells, which in the first two embeddings allude to being a differentiation checkpoint for the OPCs and neuroblasts. However, this observation disappears once the cell cycle genes are removed, where instead the nIPC cells have a transcriptional profile that agrees with developing cells in both glia and neuronal lineages. This reveals that the nIPC cells do not form an intermediate cell type per se, but rather cluster together due to the significant transcriptional change attributed to the rapid cell division during development. Instead, a seperate factor active at the right-hand site of the astrocyte cluster (figure 2C) suggests that there exists an intermediate checkpoint between the astrocytes and OPCs. This factor is characterized by the highly variable *Fabp7*, a regulator of both astrocytes as well as OPCs (14, 15). Further effect of the cell cycle can be observed in the small set of protruding cells in the OPC 2 cluster as it is no longer present in the last embedding. Indeed, oligodendrocytes in the developing brain have previously been shown to enter the cell cycle after reaching a more mature state (16).

**Fig. 2.**
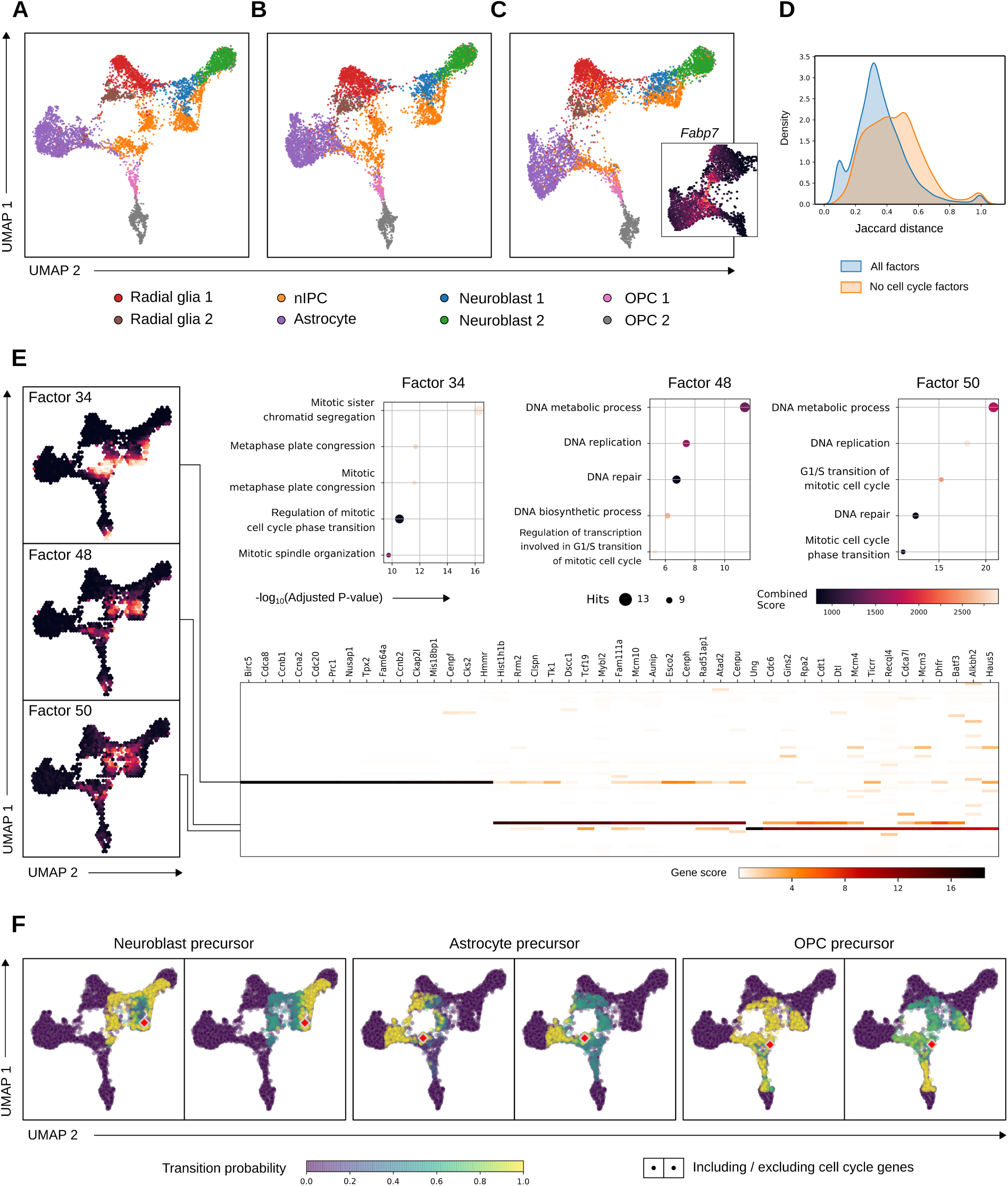
Cell cycle as a biological confounder in the developing mouse hippocampus. **A**. UMAP embedding of developing mouse hippocampus cells annotated by the cell types identified by Manno et al (12). OPC, oligodendrocyte precursor cell; nIPC, neuronal intermediate progenitor cell.**B, C** The reconstructed embeddings using the top 50 genes of each factor including and excluding the cell cycle factors identified by scHPF. **D** The distribution of Jaccard distances between the nodes of the embedding’s knn-graph and that of the embedding in A. **E** Cell cycle factors (34, 48, 50) found by scHPF identified by their Gene Ontology annotations. The cell scores are shown in the embeddings on the left. The heatmap shows the gene scores of the top 10 genes of these factors. Gene Ontology annotations are shown on top. **F** Transition probabilities including (left) and excluding (right) the cell cycle genes for three selected cells. The latter shows a clearer transition affinity to one of three cell types (rows).

### RNA velocity is influenced by confounding factors

A scRNA-seq dataset is a static snapshot of the transcriptional state of a cell and therefore does not reveal the regulation of a gene, i.e. if it is currently up- or downregulated or in a steady state of transcription. However by comparing the fraction of spliced to unspliced counts, RNA velocity allows the quantification of a gene’s regulatory state, effectively extracting a time component from the static snapshot (12). This information was used to further explore the effects of the cell cycle dynamics by calculating the cell-to-cell transition probabilities with and without the previously identified cell cycle genes. Figure 2F shows three examples of nIPC cells that show a strong transition probability to other nIPC cells when cell cycle genes are included in the calculation, with no clear commitment to any one cell fate. This implies that the transition between the stages of the cell cycle dominate, or at least contribute significantly to the aggregated velocities. However, when excluding the velocities of the cell cycle genes in the calculation of the cell transition probabilities, a much clearer picture can be established of the differentiation, as the probabilities becomes more concentrated at the different cell type clusters.

This shows that the intermediate nIPC cells already have a clear commitment to a certain cell type, an observation that is missed when selecting genes exclusively on dominant signal rather than biological contribution as showcased before in figure 2A. Previous results from the data source (12) further examplify this, as no transition from radial glia to the nIPC cluster can be observed in the velocity field.

### Matrix factorization captures regulators of cell differentiation

The second dataset from La Manno et al. is of developing glutamatergic neurons in the human forebrain. It follows a linear path from the radial glia progenitor cells to the mature neurons through a sequence of differentiation checkpoints. These checkpoints were captured in six out of 15 factors generated with scHPF and are shown sequentially in figure 3. For each of these factors, the cell scores on the embedding are shown in addition to the expression dynamics of the top 10 most contributing genes recovered by pseudotime inference using Ouija. Factor genes are largely preserved even with variable numbers of K (see Supplementary Figure S2). The first factor, factor 8, captures the radial glia cells identified by the highest scoring gene, the homeobox transcription factor *HOPX*, a well known marker gene for radial glia (12, 17). Factor 6 follows immediately after and is marked by an early deactivation of the *KLF5* gene which belongs to the Kruppel-like family of transcription factors. *KLF5* and other KLF genes are known repressors of neurite growth, and their down-regulation are linked to cell cycle arrest and neuronal development (18, 19). The later suppression of Vimentin (*VIM*) in the same factor, which is a highly variable gene (figure S1) and known marker of gliogenesis, indicates that this might be a commitment point of the progenitor cells to either neuronal or astrocytic cell fates. Factor 7 has *EOMES* as the highest scoring gene which is a well known transcription factor in early neuroblasts that regulates neurogenesis (12). Though no direct link can be found between *EPB41L4A* and neurogenesis, the early-activated *SDC2* gene has a known role in regulating axon morphology in developing neurons in mice (20). This function is continued in factor 3 where GO terms such as *regulation of dendritic spine morphogenesis* and *regulation of cell morphogenesis involved in differentiation* are highly enriched. Of note in this factor is *LAMC1*, a target gene of miR-124, a microRNA that has an important regulatory function in differentiation and maturation of neuroblasts (21, 22). The exitatory effects of glutamatergic neurons are already specified by the *GPR39* gene in factor 3 and is further regulated by the two synaptotagmins *SYT4* and *SYT1* (figure S1) in factor 1. The activity of synaptotagmins are indicative of synaptic integration, the final stage of neurogenesis (21, 23, 24). The highly variable *LMO3* gene in the final factor is a co-factor that physically interacts with other regulatory proteins to form transcriptional regulators in the developing brain (25, 26). Together these results show that genes captured with matrix factorization have important regulatory roles in the timing of neuronal differentiation.

**Fig. 3.**
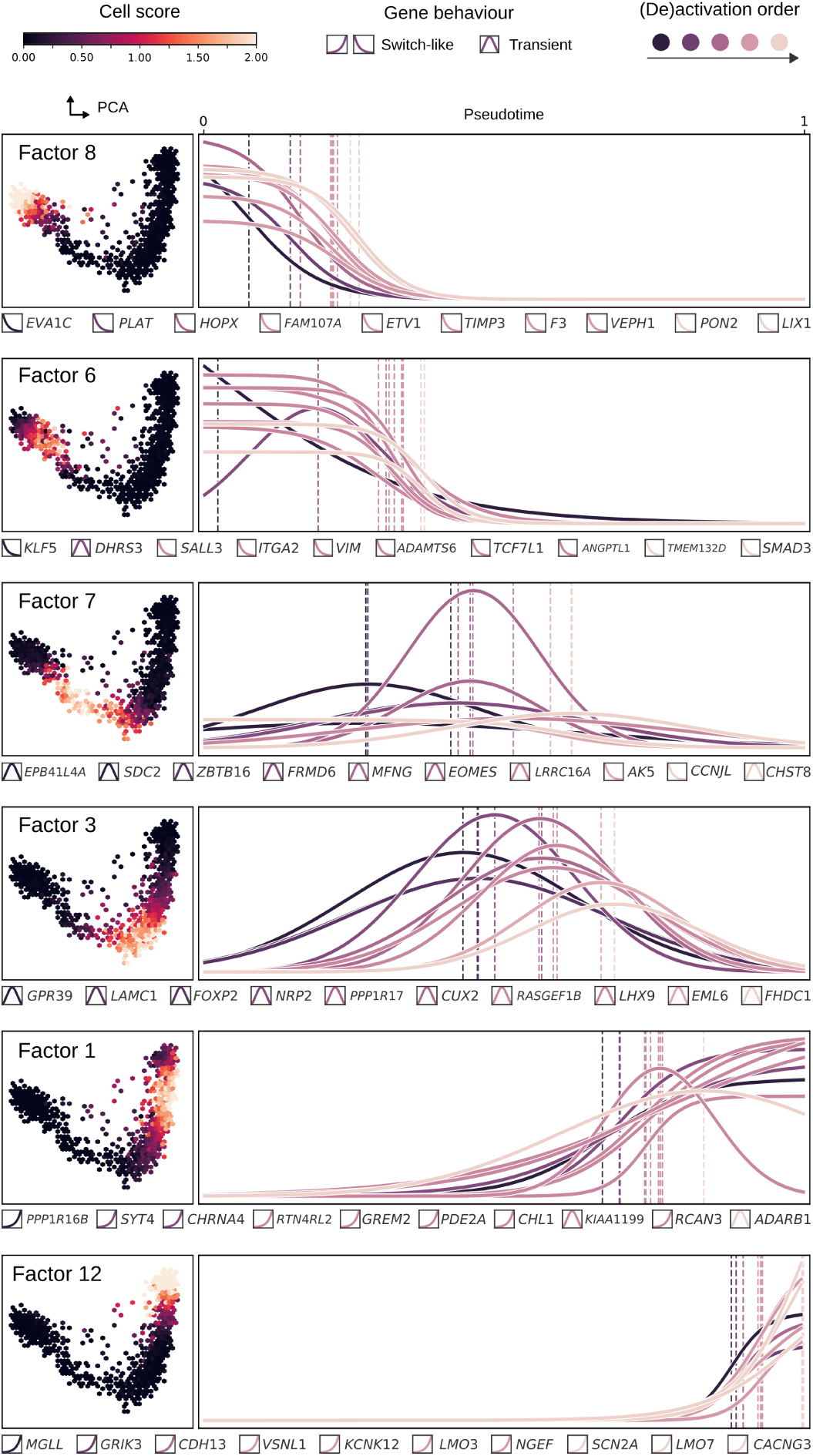
Gene expression dynamics of cell type and differentiation checkpoint factors. Each row shows for each factor the cell scores plotted on the first two principal components. On the right are the Ouija fitted curves of the top 10 factor genes. Each vertical line corresponds to the activation/deactivation time for genes with switch-like behavior, and peak time for transiently activated genes. Curve and line color indicate order of activation and deactivation. Gene names with corresponding expression behaviour type is shown below each plot.

### Pseudotime curve fitting ignored biological confounds

We compared the pseudotime assignment obtained using Ouija on the factor genes (shown in figure 3), with the pseudotimes identified by the data source, where a principle curve was fit on the first four principal components. Figure 4A shows the fitted principal curve while figure 4B shows the alignment of the cells based on the two pseudotime assigned by the two methods. A disagreement is visible in the first half of the trajectory with a delay in the radial glia cluster (blue) by Ouija. This delay is propagated until the midpoint is reached which corresponds to the start of factor 3. We hypothesise the cause of the delay to be due to confounding variability within the radial glia cluster, most likely related to cell cycle effects. As the true labels are unknown, we resort to working on the basic assumption that early down-regulated genes (e.g. those found in factors 8 and 6) are to be negatively correlated with pseudotime. A higher correlation value would therefore indicate a better ordering of cells along the pseudotime axis. Figure 4C shows the Pearson correlations that support the idea of a necessary delay within the radial glia cluster as captured by our approach. Scatterplot of the downregulted genes as a function of both pseudotime assignments are shown in supplementary figure S5.

**Fig. 4.**
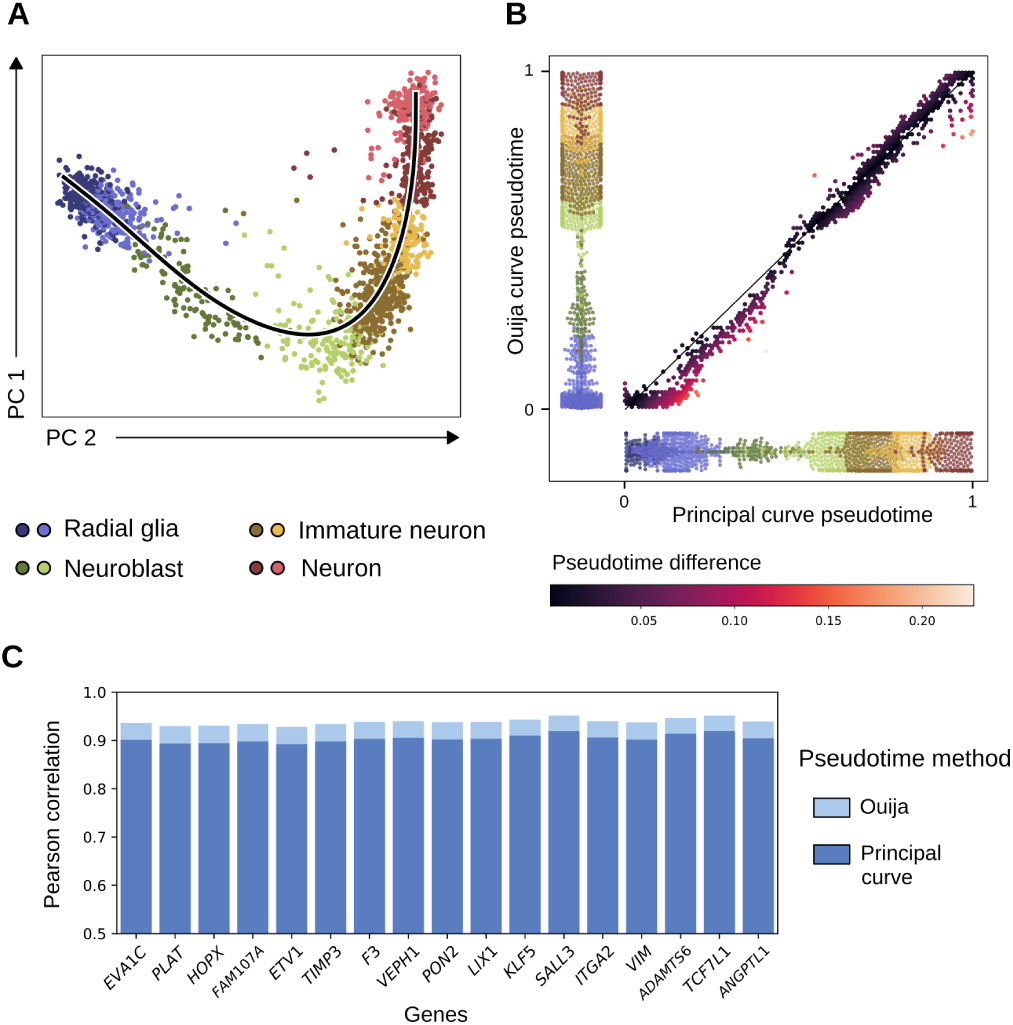
A Cells of developing human forebrain projected on the first two principal components. Black line indicates the fitted principal curve. Colors indicate clusters. **B** Cells plotted based on the pseudotime of the principal curve (x-axis) and Ouija (y-axis). Cells are colored by difference in pseudotime between the two methods. Shown behind each axis are the distribution of the cells on the basis of the pseudotime assignments, colored according to their cluster. **C** Pearson correlation between the normalized expression values and the pseudotimes of the downregulated genes of factors 8 and 6 (see figure 3).

### Matrix factorization decouples cell type and cell states

Cells within each cell type cluster can be further divided into cell states. One such example are the radial glia, which differentiate to produce both neurons and astrocytes, but have been shown to switch their preference from the former to the latter at a later stage (27–29). Figure 5A shows how matrix factorization has captured two subpoplations within the radial glia cluster, one present exclusively in postnatal day 0 and the other in postnatal day 5. The latter also shows a stark increase in astrocytes supporting the observation of two states of preferences. Another example of different cell states within the same cell type cluster are the astrocytes themselves. Among other differences, astrocytes are long known to specialize as GFAP-positive protoplasmic and GFAP-negative fibrous astrocytes (27, 30, 31). The factor shown in figure 5B identified such a subpopulation within the astrocyte cluster with a high specificity for GFAP, which might reflect the protoplasmic astrocyte state. More examples are found in the second dataset where multiple factors are found that overlap with the cell type and checkpoint factors identified in figure 3 (figure S1).

**Fig. 5.**
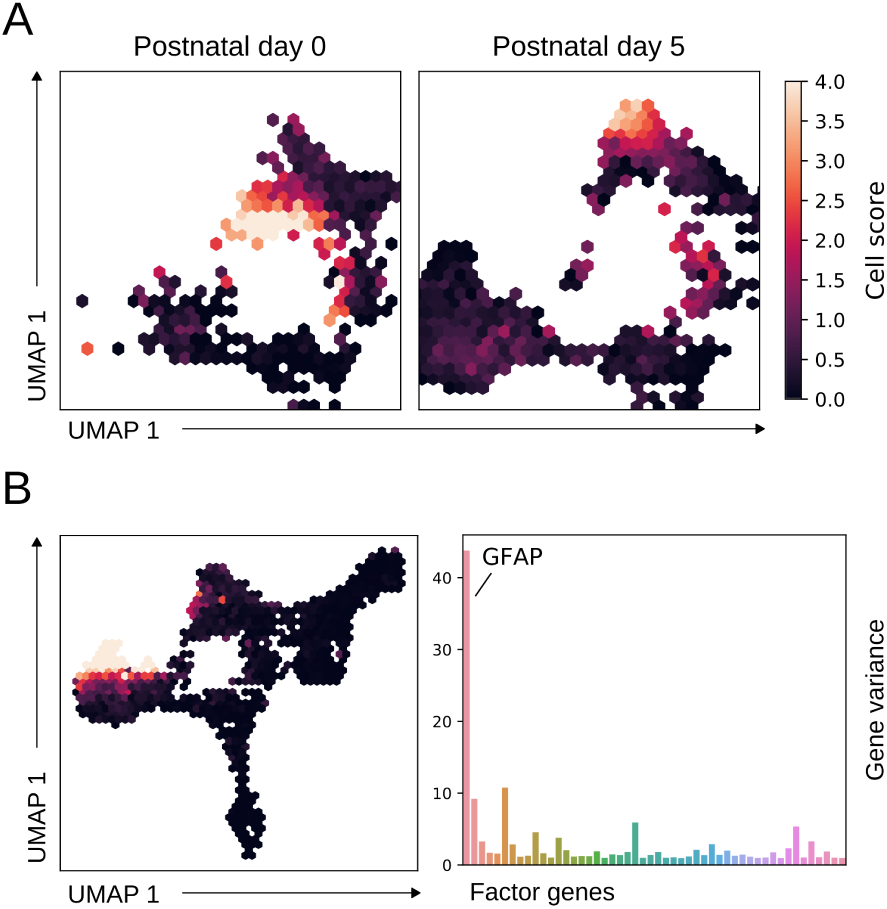
Cell states within cell types. **A** Cell subsets from the hippocampus dataset shown in figure 2A plotted based on the timepoint of extraction. Plots are truncated to show mainly radial glia and astrocytes. Radial glia first prioritize production of neurons, hence no astrocytes are visible at postnatal day 0. However, prduction is switched to astrocytes afterwards. Matrix factorization is able to capture these two different states of the radial glia. **B** Factor active in a subset of astrocyte cells. The high specificity of *GFAP* in the factor alludes to it being the protoplasmic subtype.

## Discussion

We have made an attempt at understanding the complex process of cell differentiation by directly modelling regulators of selected transcriptional programs presumed to be cell type-specific. This approach tackles the question of what biological sources of variation are relevant to developing cells. We followed a “bottom-up” approach where a small subset of core marker genes is utilized to represent cell identity. Alternatively, one can follow a top down approach such that all biological processes active in the cells are annotated and subsequently stripped away based on their perceived relevance. An example of this is of Buettner et al. (2), where the same problem formulation of confounding cell states was tackled. There, a latent variable model was developed that factorizes the expression matrix into a set of factors determined by a database of pathways and another set of unannotated factors. However, their analysis was limited to removing technical rather than biological confounding sources of variation. Nonetheless we argue that while both approaches require a fair amount of domain knowledge and manual interpretation of gene sets, a bottom-up approach alleviates this by a large margin while still adhering to a concrete definition of cell identity.

### Dimensionality reduction in pseudotime inference

The end goal of our methodology is to improve pseudotime inference rather than to develop a new algorithm, and to shift the discussion towards the use of biology motivated tooling and interpretation. This is in response to most tools developed for reconstructing trajectories from single cell data, namely that they adhere to the pipeline of dimensionality reduction followed by trajectory modelling, by either fitting a curve in the resulting embedding or by finding a path in the embedding’s neighbour graph (10, 32). While it is known that dimensionality reduction leads to a loss of interpretability, other relevant concerns can be raised as well. Firstly, the issue of the frequently used PCA method in count based data such as scRNA-seq was recently addressed by (amongst others) Townes et al. (33), where the authors show that there exists an implicit assumption of normality of the features. The authors show that this results in distorted components as scRNA-seq data violates this assumption. Secondly, non-linear dimensionality reduction methods such as t-SNE and UMAP can exaggerate the distances between cell clusters with large transcriptional differences, which can lead to disconnected embeddings (34). This problem can be amplified with stringent variance-based gene filtering as developing cells are identified by more subtle differences in gene expression. Finally, it has been shown that complex high-dimensional structures found in scRNA-seq data cannot be fully preserved in a small number of dimensions, and can therefore miss important variation or lead to distorted embeddings (35). Our methodology is unaffected by these issues as dimensionality reduction is not a prerequisite for matrix factorization and gene expression modelling. However it must be noted that the use of gene expression smoothing used in the RNA velocity pipeline, which was subsequently passed to Ouija for gene modelling, does rely on a PCA step, and may therefore be affected by possible distortions mentioned before. This is no shortcoming of Ouija as no smoothing is required per se, however we did find the model fit to improve with smoothing (supplementary Figures S3 and S4).

### Cell type and cell state

Of specific interest in this study is the definition of a cell type as a summation of its core regulators and its different states. Trajectory modelling is affected by this as cell distances can reflect cell state instead of cell types. We argue that reducing the genes to a small set of core regulators that preserve the identity of the cells, we are able to circumvent the problem of confounding cell states. Another consequence of decoupling cell type from state is that a shared state between different cell types can lead to high similarity between cells of different identities. This is exemplified in the first analysis shown in figure 2 where it was found that the existence of the nIPC cluster is a direct effect of the cell cycle on differentiating cells. A more subtle example is regarding the substantial amount of functional similarities between radial glia and astrocytes, hence their combined classification as neuroglia (31). The expression of the GFAP protein in both cell types is an example of their similarities, which can explain the close proximity observed in figure 2B between the radial glia and the aforementioned protoplasmic astrocyte subpopulation.

### Regulator genes for cell type identification

We have motivated and shown that the use of cell type regulators for pseudotime inference can be a useful alternative to curve-based fitting due to their biological relevancy. However not all transcription factors are annotated or even identified (36). This problem is circumvented in this study by taking the top factor genes, and although many have important regulatory functions as shown in the results, it is not obvious if they are directly coupled to the cell types and differentiation checkpoints. Further validation must be needed to confirm such associations. For example, knockout experiments of the putative transcription factors can be used to determine their necessity as important regulators in cell differentiation. Similarly, one can overexpress the putative regulator in pluripotent stem cells and observe any structural or regulatory similarities with the cell type under study. One might argue that as long as the genes used for pseudotime inference are faithful proxies of the differentiation process, the use of effector genes rather than regulatory genes might not influence the resulting pseudotimes significantly. A downside to this however is that the genes cannot be validated or utilized in other similar studies which hurts the interoperability and reusability of the study. Furthermore, the genes may no longer provide insight into what processes regulate cell fate decisions. Effector genes are also much larger in quantity than their regulators (1), which can lead to an uneven distribution of signals across the different cell types and checkpoints and result in distorted pseudotimes. Even when restricting the number of genes to circumvent this, one must also keep in mind that there exists a delay in activity between transcription factors and effector genes. Namely, when a pathway used during differentiation is activated in developing cells, there exists a delay between when its regulators respond to the activation signals, and the actual transcription and activity of the effector genes. This means that the snapshot provided by scRNA-seq of the transcript counts fluctate in time, which has led to the utlization of time alignment algorithms such as Dynamic Time Warping, used predominantly in the field of metabolomics (37). Current computational solutions might be the use of regulatory network inference algorithms that have emerged in quantity due to the granularity provided by scRNA-seq data (38). Another approach is provided by algorithms that predict physical interactions between proteins, as increasing evidence shows that core regulators form physical interactions (1), exemplified by the co-factor LMO3 found in factor 12 in the developing mouse forebrain. Figure S6 in the supplementary shows how the STRING service (39) is used to find many interactions between top factor genes in the last stages of glutamatergic neurogenesis.

## Conclusion

The incredible resolution provided by single cell RNA-sequencing data raises the question of what sources of variation within it are important for the study at hand. Here we have focused on the problem of cell identification, more specifically in developmental systems using pseudotime inference. We have argued and shown that confounding sources of variation, most notably the cell cycle, can distort inference of the differentiation trajectory. We have then shown that this problem can be circumvented by limiting the scope to a select subset of genes assumed to play a regulatory role in cell development and directly modelling their expression.

## Materials and Methods

### Data and preprocessing

Two single cell RNA-seq datasets with continuous cell type transitions were acquired from La Manno et al. 2018 (12): Developing glutamatergic neurons in the human forebrain, which has a well-defined linear manifold, and a complex branching dataset of the developing mouse hippocampus (Gene Expression Omnibus accession code GSE104323). Both datasets were available as raw unspliced and spliced count matrices with the corresponding gene and cell annotations. Normalization was done with Seurat 3.0’s scTransform (40). In short, a regularized negative binomial model is fit on the data to model the counts as a function of total cell size. The Pearson residuals yielded by this fit can then be treated as normalized expression values, where a positive value indicates a higher count than expected, and vice versa. Seurat uses these residuals to return a count matrix that is unbiased by cell size, which was used for pseudotime inference. However, scHPF models the genes directly as negative binomial and was therefore passed the unnormalized spliced matrix instead. Filtering of low quality genes and cells was done in correspondence to the Jupyter notebooks provided by the authors (https://github.com/velocyto-team/velocyto-notebooks): In the human forebrain dataset, genes in the unspliced matrix with a total count lower than 25 or a minimum cell count less than 20 were removed, and in the spliced matrix the thresholds were set at 30 and 20 respectively. The 1,720 cells in the dataset were not filtered. In the mouse hippocampus dataset, the same gene threshold was set for the unspliced matrix while the thresholds were increased to 40 and 30 respectively for the spliced matrix. Cells with a total gene count lower than the 0.4 precentile were also filtered out. In addition, neuronal clusters identified by the source (Subiculum, CA1, CA3, CA2/4, Granule) were removed, leaving a total of 6,673 out of 18,213 cells. Variant genes in both datasets were selected on the basis of residual variance provided by the model fit rather than the mean-variance association.

### Cell cycle effect analysis

The branching mouse hippocampus dataset was truncated by removing the neuronal specification branches identified by the original paper, which included everything beyond the neuroblast clusters. A UMAP embedding was created with parameters *n*_*neighbour* = 30, *min*_*dist* = 0.1, and *metric* = *correlation*. scHPF was run with *K* = 60 (number of factors) and the resulting factors were annotated by Gene Ontology (GO) enrichment of the top scoring factor genes using the Enrichr webservice (41). Factors with enriched GO terms within the cell cycle hierarchy were recognized as cell cycle factors (figure 2E). Two new UMAP embeddings were then calculated with the top scoring factor genes, one including and one excluding the cell cycle factors. To quantify the agreement of both embeddings with the original, the distribution of Jaccard distances between the 500 nearest neighbours of each cell were calculated (figure 2D). Results were validated using RNA velocity, a method by which the RNA splicing dynamics are used to extrapolate the expression profile at a future timepoint (12). The similarity of the extrapolated profile of a cell and that of the measured profiles of all other cells can be restated as a transition probability (Supplementary Note 1 of (12)). In summary, the transition probability between two cells *i* and *j* is found by calculating the Pearson correlation of the difference between the two cell profiles and the RNA velocity vector of cell *i*, which is then passed through an exponential kernel. Repeating this procedure for each pair of cells yields a matrix of similarity values which is then transformed into probability-like values by normalizing the rows to sum up to 1. These cell-to-cell transition probabilities were used to showcase the effect of cell cycle on the transition.

### Pseudotime linear dataset

The human glutamatergic neurogenesis dataset was factorized into 15 scHPF factors. Cell type and differentiation checkpoint factors were selected on the basis of known marker genes and enriched GO terms linked to neuronal differentiation such as *dendrite extension* and *cell morphogenesis in differentiation*. The top 10 factor genes were selected and subsequently passed to Ouija for pseudotime inference. However, kNN-smoothed expression values were passed instead of the log-transformed counts as this leads to a better model fit (figure S3). A principal curve was also fitted on the first four principal components following the afore-mentioned notebook provided by the data source. Pseudotimes of the two methods were then compared by simply calculating the differences in pseudotime assignment per cell. This revealed a disagreement in pseudotime assignment between the two methods. This disagreement was evaluated by correlating the down-regulated genes with the pseudotimes under the basic presumption that these should be negatively correlated, i.e. down-regulated genes decreases monotonically in expression as a function of pseudotime.

### Data availability

Both datasets are acquaired from La Manno et al. 2018 (12). The developing mouse hippocampus dataset is available in the Gene Expression Omnibus with accession code GSE104323. Analysis scripts can be found as a Snakemake pipeline in https://github.com/mochar/cell_variation.

## Supplementary Information

**Fig. S1.**
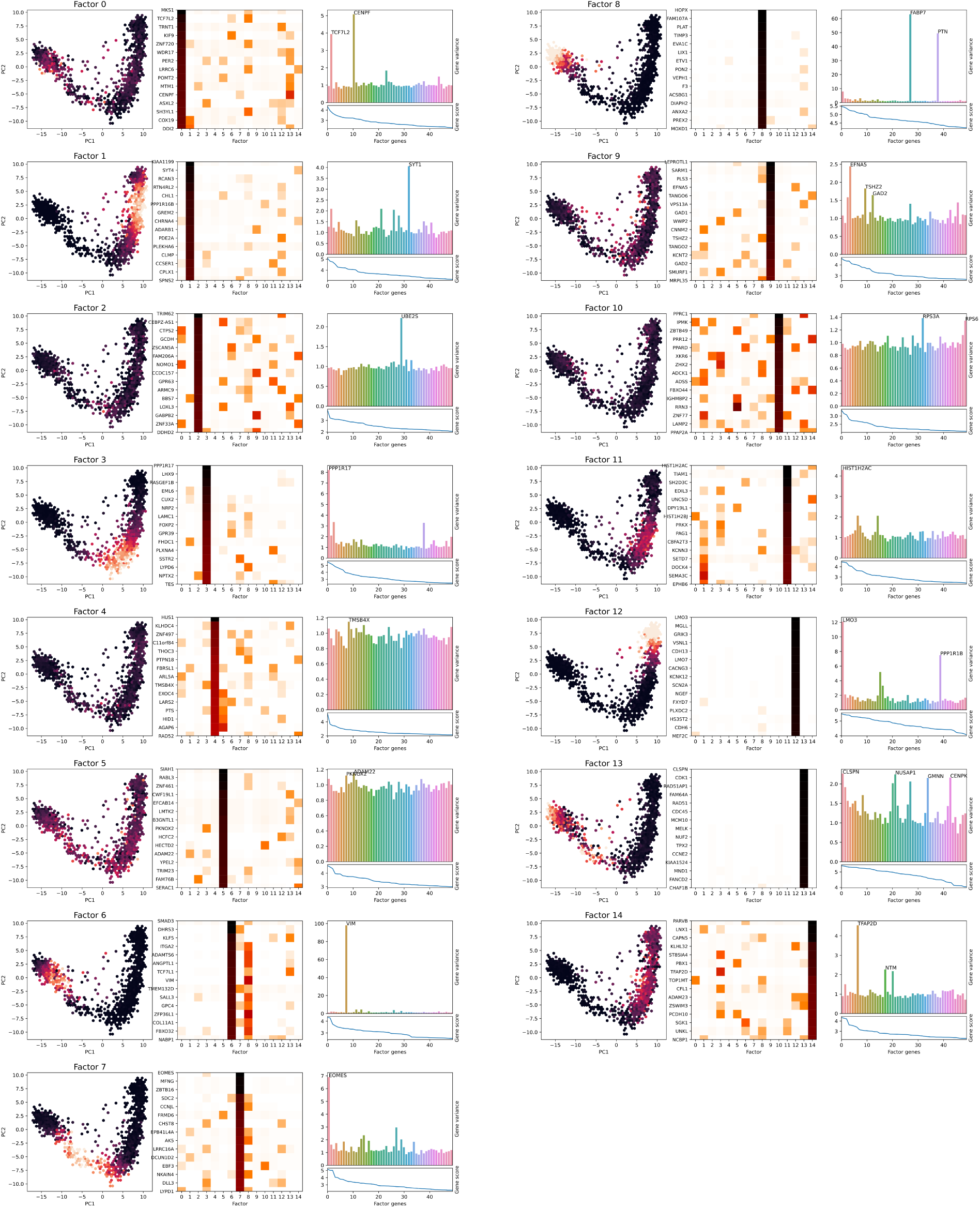
Factors identified in the human neurogenesis dataset. Shown are for each factor: the cell scores in the UMAP embedding (left), the gene scores of the top 15 factor genes (middle), and the gene variance sorted by factor score of the top 50 factor genes (right).

**Fig. S2.**
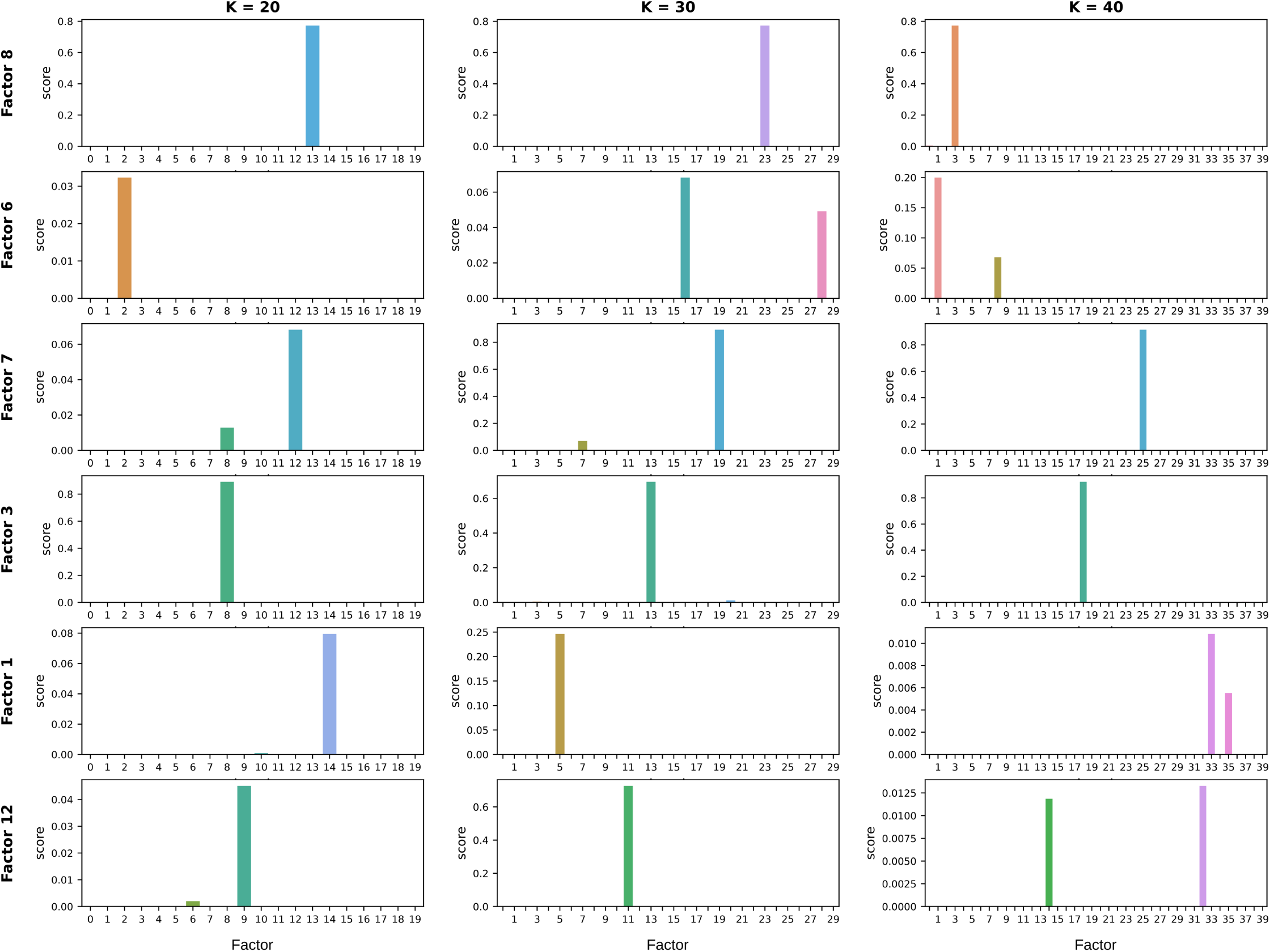
Factor perseverance across different K values for the factors shown in figure S1. The rank-biased overlap score of the top 1000 genes were calculated for all factors generated with K = 20, 30 and 40. Original factors were calculated with K = 15.

**Fig. S3.**
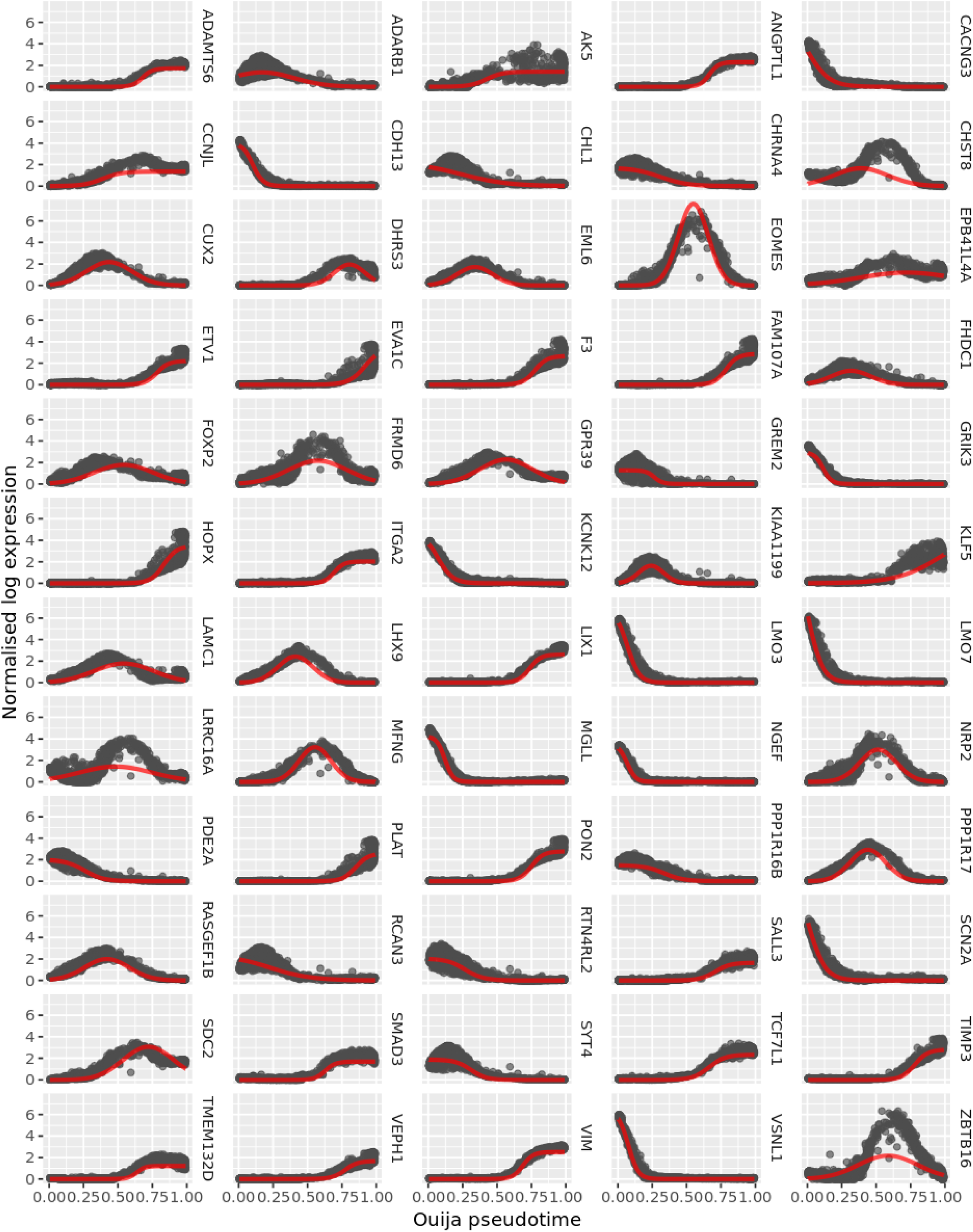
Ouija gene expression modelling fits of the top 10 genes of the factors shown in figure 3.

**Fig. S4.**
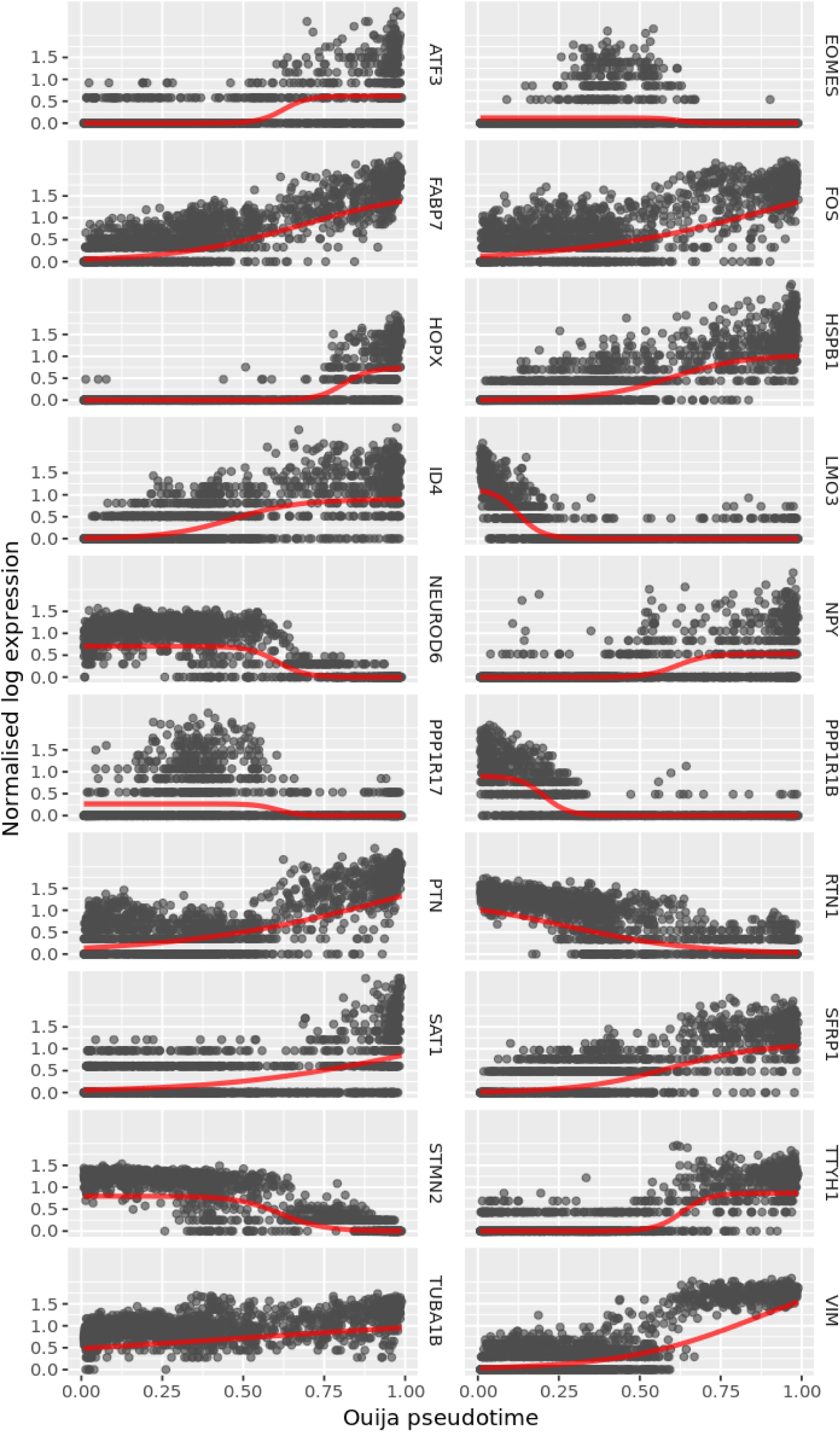
Old Ouija run without gene smoothing.

**Fig. S5.**
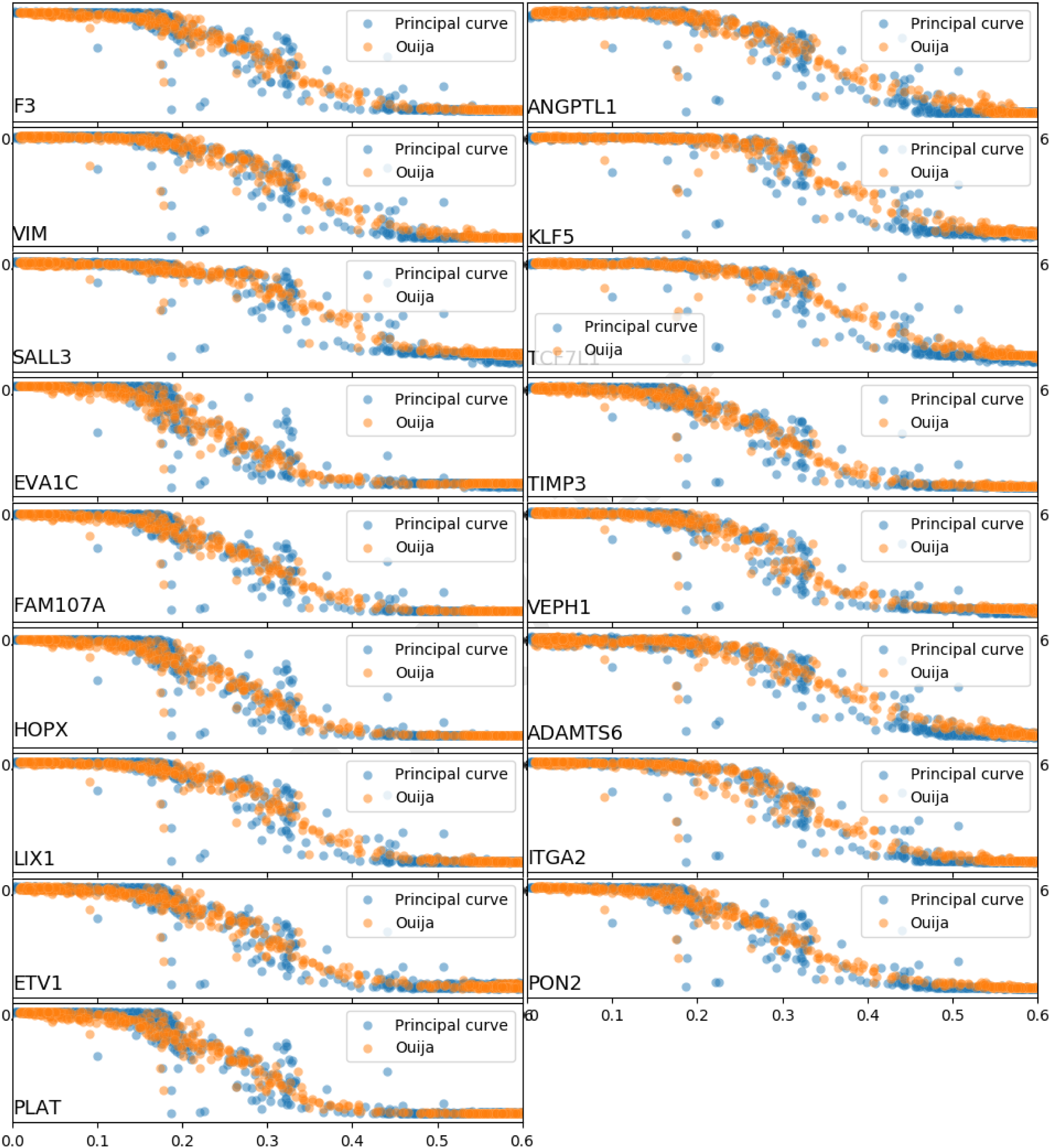
Gene expression of early down-regulated genes as a function of pseudotime determined by principle curve fitting and Ouija fitting.

**Fig. S6.**
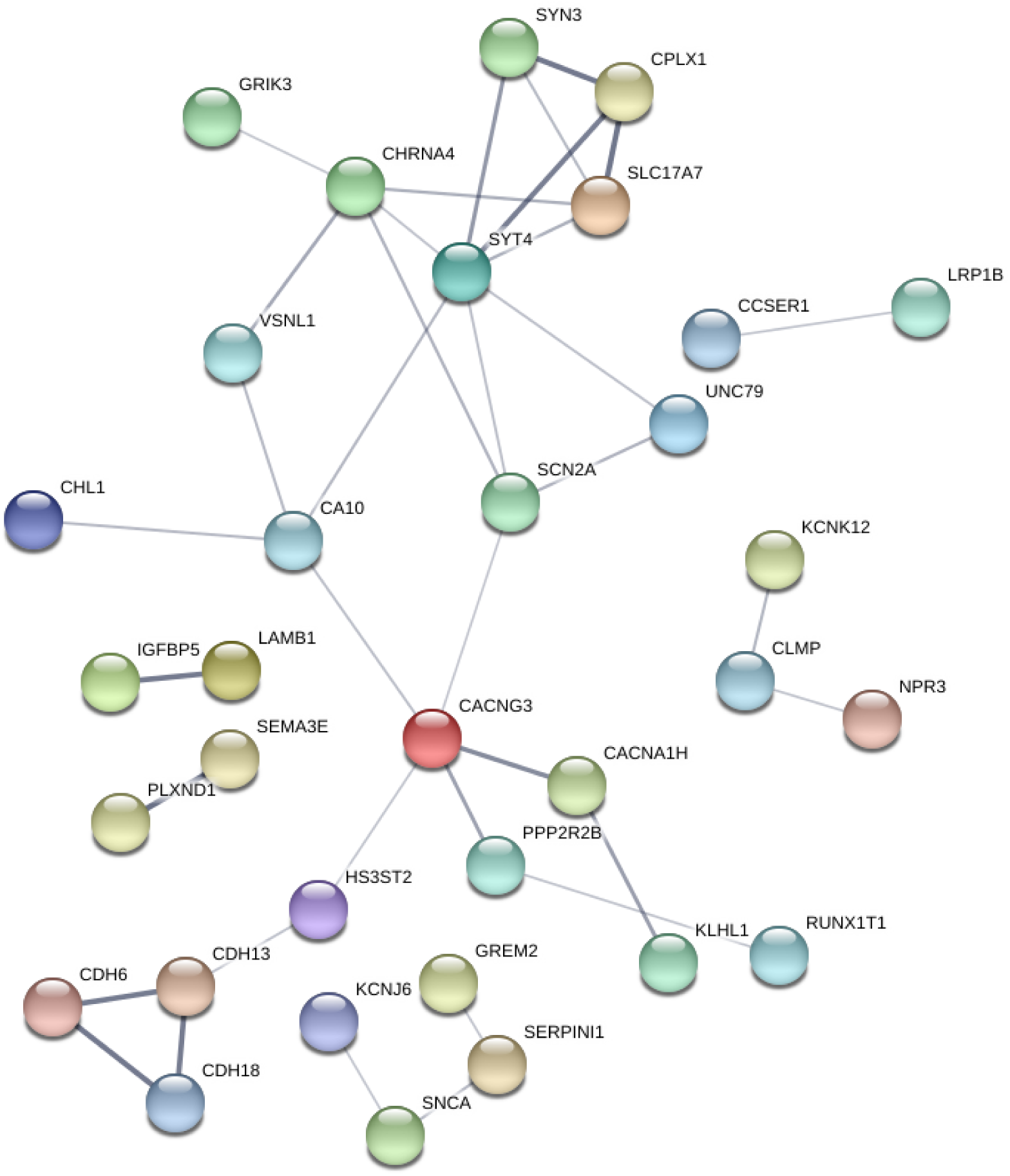
Protein-protein interactions predicted by STRING. Inputted were the top 30 genes of the last two factors shown in figure 3

## References

1. D Arendt, et al., The origin and evolution of cell types. Nat. Rev. Genet. 17, 744–757 (2016).

2. F Buettner, et al., Computational analysis of cell-to-cell heterogeneity in single-cell rna-sequencing data reveals hidden subpopulations of cells. Nat. Biotechnol. 33, 155–160 (2015).

3. DH Nguyen, RG Jaszczak, DJ Laird, Chapter Five - Heterogeneity of primordial germ cells, The Immortal Germline, ed. R Lehmann. (Academic Press) Vol. 135, p. 155–201 (2019).

4. RL Davis, H Weintraub, AB Lassar, Expression of a single transfected cdna converts fibrob-lasts to myoblasts. Cell 51, 987–1000 (1987).

5. T Graf, T Enver, Forcing cells to change lineages. Nature 462, 587–594 (2009).

6. K Takahashi, S Yamanaka, Induction of pluripotent stem cells from mouse embryonic and adult fibroblast cultures by defined factors. Cell 126, 663–676 (2006).

7. B Xia, I Yanai, A periodic table of cell types. Development 146, dev169854 (2019).

8. JL McFaline-Figueroa, et al., A pooled single-cell genetic screen identifies regulatory checkpoints in the continuum of the epithelial-to-mesenchymal transition. Nat. Genet. 51, 1389–1398 (2019).

9. KR Campbell, C Yau, A descriptive marker gene approach to single-cell pseudotime inference. Bioinformatics, 10 (2018).

10. W Saelens, R Cannoodt, H Todorov, Y Saeys, A comparison of single-cell trajectory inference methods: towards more accurate and robust tools. Nat. Biotechnol., 49 (year?).

11. HM Levitin, et al., De novo gene signature identification from single-cell rna-seq with hierarchical poisson factorization. Mol. Syst. Biol. 15, e8557 (2019).

12. G La Manno, et al., Rna velocity of single cells. Nature 560, 494–498 (2018).

13. P Malatesta, et al., Neuronal or glial progeny: Regional differences in radial glia fate. Neuron 37, 751–764 (2003).

14. K Sharifi, et al., Fabp7 expression in normal and stab-injured brain cortex and its role in astrocyte proliferation. Histochem. Cell Biol. 136, 501 (2011).

15. K Sharifi, et al., Differential expression and regulatory roles of fabp5 and fabp7 in oligodendrocyte lineage cells. Cell Tissue Res. 354, 683–695 (2013).

16. SA Goldman, NJ Kuypers, How to make an oligodendrocyte. Development 142, 3983–3995 (2015).

17. H Hochgerner, A Zeisel, P Lönnerberg, S Linnarsson, Conserved properties of dentate gyrus neurogenesis across postnatal development revealed by single-cell rna sequencing. Nat. Neurosci. 21, 290–299 (2018).

18. S Qin, CL Zhang, Role of krüppel-like factor 4 in neurogenesis and radial neuronal migration in the developing cerebral cortex. Mol. Cell. Biol. 32, 4297–4305 (2012).

19. DL Moore, A Apara, JL Goldberg, Kruppel-like transcription factors in the nervous system: Novel players in neurite outgrowth and axon regeneration. Mol. cellular neurosciences 47, 233–243 (2011).

20. B Barak, N Feldman, E Okun, Toll-like receptors as developmental tools that regulate neurogenesis during development: an update. Front. Neurosci. 8 (2014).

21. P Bielefeld, C Mooney, DC Henshall, CP Fitzsimons, mirna-mediated regulation of adult hippocampal neurogenesis; implications for epilepsy. Brain Plast. 3, 43–59 (2017).

22. MF Lang, Y Shi, Dynamic roles of micrornas in neurogenesis. Front. Neurosci. 6 (2012).

23. G Tocco, et al., Two synaptotagmin genes, syt1 and syt4, are differentially regulated in adult brain and during postnatal development following kainic acid-induced seizures. Mol. Brain Res. 40, 229–239 (1996).

24. B Ullrich, et al., Functional properties of multiple synaptotagmins in brain. Neuron 13, 1281–1291 (1994).

25. A Abellán, E Desfilis, L Medina, Combinatorial expression of lef1, lhx2, lhx5, lhx9, lmo3, lmo4, and prox1 helps to identify comparable subdivisions in the developing hippocampal formation of mouse and chicken. Front. Neuroanat. 8 (2014).

26. M Sang, et al., Lim-domain-only proteins: multifunctional nuclear transcription coregulators that interacts with diverse proteins. Mol. Biol. Reports 41, 1067–1073 (2014).

27. H Tabata, Diverse subtypes of astrocytes and their development during corticogenesis. Front. Neurosci. 9 (2015).

28. AV Molofsky, B Deneen, Astrocyte development: A guide for the perplexed. Glia 63, 1320–1329 (2015).

29. R Beattie, S Hippenmeyer, Mechanisms of radial glia progenitor cell lineage progression. Febs Lett. 591, 3993–4008 (2017).

30. OA Bayraktar, LC Fuentealba, A Alvarez-Buylla, DH Rowitch, Astrocyte development and heterogeneity. Cold Spring Harb. Perspectives Biol. 7, a020362 (2015).

31. T Mori, A Buffo, M Götz, The Novel Roles of Glial Cells Revisited: The Contribution of Radial Glia and Astrocytes to Neurogenesis. (Elsevier) Vol. 69, p. 67–99 (2005).

32. R Cannoodt, W Saelens, Y Saeys, Computational methods for trajectory inference from single-cell transcriptomics. Eur. J. Immunol. 46, 2496–2506 (2016).

33. FW Townes, SC Hicks, MJ Aryee, RA Irizarry, Feature selection and dimension reduction for single cell rna-seq based on a multinomial model. bioRxiv, 574574 (2019).

34. A Konstorum, N Jekel, E Vidal, R Laubenbacher, Comparative analysis of linear and nonlinear dimension reduction techniques on mass cytometry data. bioRxiv, 273862 (2018).

35. SM Cooley, T Hamilton, EJ Deeds, JCJ Ray, A novel metric reveals previously unrecognized distortion in dimensionality reduction of scrna-seq data. bioRxiv, 689851 (2019).

36. SA Lambert, et al., The human transcription factors. Cell 172, 650–665 (2018).

37. C Christin, et al., Time alignment algorithms based on selected mass traces for complex lc-ms data. J. Proteome Res. 9, 1483–1495 (2010).

38. H Todorov, R Cannoodt, W Saelens, Y Saeys, Network inference from single-cell transcriptomic data. Methods Mol. Biol. (Clifton, N.J.) 1883, 235–249 (2019).

39. D Szklarczyk, et al., String v11: protein-protein association networks with increased coverage, supporting functional discovery in genome-wide experimental datasets. Nucleic Acids Res. 47, D607–D613 (2019).

40. C Hafemeister, R Satija, Normalization and variance stabilization of single-cell rna-seq data using regularized negative binomial regression. bioRxiv, 576827 (2019).

41. MV Kuleshov, et al., Enrichr: a comprehensive gene set enrichment analysis web server 2016 update. Nucleic Acids Res. 44, W90–97 (2016).

